# Amino acid availability acts as a metabolic rheostat to determine the magnitude of ILC2 responses

**DOI:** 10.1101/2022.06.22.497162

**Authors:** Suzanne H. Hodge, Maria Z. Krauss, Irem Kaymak, James King, Andrew J.M. Howden, Gordana Panic, Richard K. Grencis, Jonathan R. Swann, Linda V. Sinclair, Matthew R. Hepworth

**Affiliations:** Lydia Becker Institute of Immunology and Inflammation, University of Manchester, M13 9PL, United Kingdom; Division of Infection, Immunity and Respiratory Medicine, School of Biological Sciences, Faculty of Biology, Medicine and Health, Manchester Academic Health Science Centre, University of Manchester, M13 9PL, United Kingdom; Cell Signalling and Immunology Division, School of Life Sciences, University of Dundee, Dundee, DD1 5EH, United Kingdom; Division of Integrative Systems Medicine and Digestive Diseases, Imperial College London, South Kensington, SW7 2AZ, United Kingdom; School of Human Development and Health, Faculty of Medicine, University of Southampton, S016 6YD, United Kingdom

**Keywords:** Innate lymphoid cells, ILC2, metabolism, helminth, solute carriers

## Abstract

Group 2 innate lymphoid cells (ILC2) are functionally poised, tissue-resident lymphocytes that respond rapidly to damage and infection at mucosal barrier sites. ILC2 reside within complex microenvironments where they are subject to cues from the diet, commensal microbiota and invading pathogens – most notably helminths. Emerging evidence suggests ILC2 are acutely sensitive not only to canonical activating signals, but also perturbations in nutrient and metabolite availability. In the context of helminth infection, we identify amino acid availability as a nutritional cue in regulating ILC2 responses. ILC2 were found to be uniquely pre-primed to import amino acids via the large neutral amino acid transporters *Slc7a5* and *Slc7a8*. Cell-intrinsic deletion of these transporters impaired ILC2 expansion, but not cytokine production, in part via tuning of mTOR activation. These findings implicate the import of amino acids as a metabolic requisite for optimal ILC2 responses, and further highlight nutritional cues as critical regulators of innate immune responses within mucosal barrier tissues.

## Introduction

Type 2 immune responses are specialised to induce effector mechanisms that mediate protective immunity to large extracellular helminth parasites that invade and inhabit mucosal barrier tissues (*1, 2*). Indeed, helminth infections have been postulated to be the major evolutionary driver of the type 2 immune system, although the precise factors that regulate the magnitude and quality of type 2 immune cell responses remain incompletely defined. Chronic helminth infections are associated with significant morbidity – including malnutrition potentially due to competition with the host for metabolic resources, which can have potent immunomodulatory consequences (*3, 4*). Indeed, an emerging body of evidence suggests the mammalian immune system is primed to sense nutrients and metabolites derived directly from the diet or produced by the commensal microbiota or pathogenic organisms (*5, 6*). Moreover, gastrointestinal helminth infections are associated with alterations in both the microbiota and dietary nutrient availability (*3, 4, 7-9*).

Group 2 innate lymphoid cells (ILC2) are transcriptionally and functionally poised effector immune cells found primarily at mucosal barrier sites, and which respond rapidly during the early phases of helminth infection by robustly producing the effector cytokines Interleukin (IL)-5 and IL-13 (*10, 11*). Alarmin signals including IL-25, IL-33 and thymic stromal lymphopoietin (TSLP) released by non-hematopoietic cells in response to tissue damage act in concert with cues from tissue-resident neurons and glial cells to induce rapid proliferation and expansion of ILC2, and induce protective responses such as eosinophilia, goblet cell hyperplasia, epithelial cell extrusion and smooth muscle hypercontractility (*10, 11*). In addition, it is increasingly appreciated that ILC2 sense and respond to changes in the abundance and availability of dietary and microbially derived metabolites including Vitamin A-derived retinoic acid (*12*), aryl-hydrocarbon receptor (Ahr) ligands (*13*), short chain fatty acids (*14*) and succinate (*15, 16*) – suggesting ILC2 are poised to sense not only tissue-associated danger signals but also the broader metabolic milieu of mucosal tissues (*17*).

In addition, micronutrients are key determinants of immune effector function through their capacity to provide fundamental substrates for production of the energy and biomass needed to fuel proliferation and protein translation (*5, 6*). Indeed, the ability of ILC2 to mount an effective and appropriate response to challenge has been shown to be dependent upon the ability to appropriately engage cell-intrinsic metabolic pathways to catabolise glucose, fatty acids and arginine (*18-20*). Despite these advances it remains unclear whether changes in the availability of metabolites occur during infection that may determine the quality and magnitude of ILC2 responses. Moreover, the precise nature of metabolic cues that modulate ILC2 responses and underpin their rapid and innate effector functions remain incompletely defined.

Here we identify amino acid availability as a critical rheostat of ILC2 responses. Strikingly – and unlike other steady state tissue resident immune cells – ILC2 were found to express multiple solute carrier-encoded transporters that act to ensure ILC2 are pre-poised to take up essential amino acids from the environment. Notably, absence of these transporters impacted the ability of ILC2 to proliferate, but not produce effector cytokines. This was found to be in part through their ability to tune ILC2 metabolic fitness and mTOR pathway activation. Together these findings suggest that ILC2 are metabolically primed to facilitate rapid expansion following activation by alarmins or in the context of helminth infection.

## Results and Discussion

### Amino acid availability impacts type 2 immunity during helminth infection

To identify environmental and metabolic cues that could impact innate type 2 immune responses in the context of helminth infection we infected mice with *Nippostrongylus brasiliensis* for 7 days and performed unbiased metabolomic analysis on feces of infected mice and control animals (Figure 1A). Using this approach we identified a total of 32 unique molecules, of which 13 were found to differ significantly (p<0.05). We identified changes in the relative abundance of several metabolites following infection, including a relative decrease in glucose and increase in lactate in the feces of infected mice, whereas no consistent differences were observed in the abundance of common microbial metabolites such as short chain fatty acids were detected (Fig. S1A). Notably, we detected increases in the relative abundance of a number of amino acids following infection including alanine, valine, leucine and isoleucine, among others. Many of the amino acids found to be increased are “essential” amino acids that cannot be synthesised by mammalian cells and instead must be acquired from dietary and environmental sources (Figure 1A+B, Fig. S1B). Comparable analysis of mice infected with other small intestinal dwelling helminths, specifically *Heligmosomoides polygyrus* and *Trichinella spiralis*, yielded similar changes in fecal amino acid abundance, albeit to different degrees (Figure 1B, Fig. S1B).

**Figure 1.**
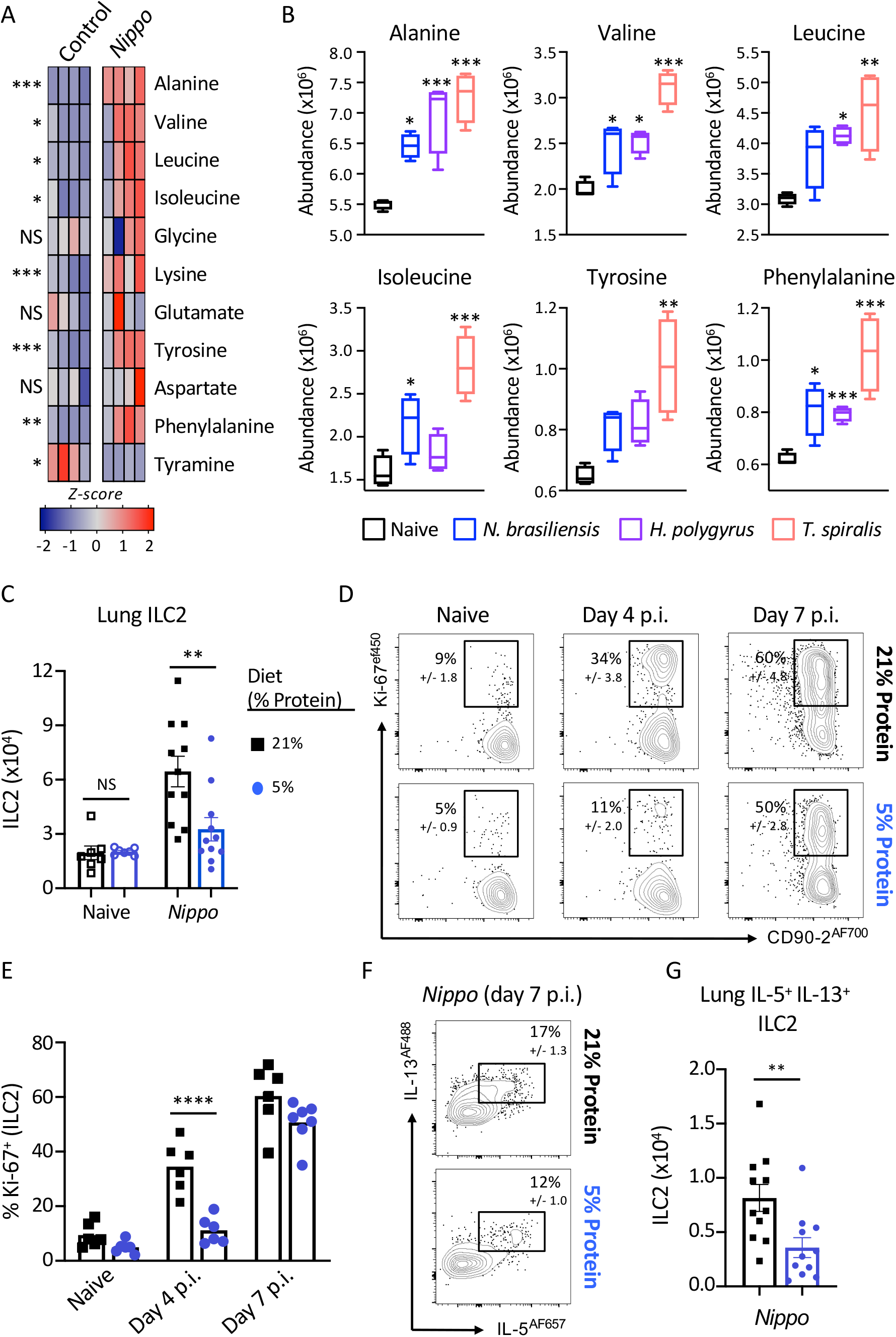
Metabolite and dietary factors influence innate type 2 response to helminth infection. A) Relative levels of fecal amino acids and amino acid-related metabolites in control and day 7 post infection *N. brasiliensis* infected C57BL/6 mice (*n*=4 mice per group, representative of two independent experiments, data shows z-scores). B) Relative abundance of selected amino acids in naïve mice or mice infected with *N. brasiliensis* (day 7 p.i. blue), *H. polygyrus* (day 7 p.i., purple) or *T. spiralis* (day 7 p.i., pink), (*n*=4 mice per group, representative of one experiment, data shows relative abundance). C) Numbers of ILC2 in naïve or *N. brasiliensis* infected (day 7 p.i.) mice, D) frequency and E) number of Ki-67^+^ ILC2 at day 4 and day 7 post *N. brasiliensis* infection, and F) frequency and G) number of IL-5 and IL-13 producing ILC2 at day 7 post *N. brasiliensis* infection in C57BL/6 mice fed a normal (21%) or low (5%) protein diet. Data shown as individual values or mean +/- SEM, * p< 0.05, ** p< 0.01, *** p< 0.001.

Given the previously reported impact of nutrient availability on ILC2 responses (*12, 18-20*), we hypothesised that an altered abundance of amino acids in the gastrointestinal tract may impact upon the quality or magnitude of a protective innate immune response during helminth infection. To test this, we fed mice a diet that was relatively low in protein (5% energy from protein), which has previously been shown to limit both tissue and systemic amino acid availability (*21*), and compared to mice fed a control diet (21% energy from protein; comparable with normal chow used in these studies). We focused our analysis on the lung – a tissue through which *N. brasiliensis* migrates during the first days of infection inducing significant tissue damage and eliciting a potent ILC2 response. Mice fed a 5% protein diet exhibited a reduced accumulation of ILC2 numbers by day 7 post-infection (Figure 1C), which was associated with a delayed proliferative response as compared to 21% protein diet-fed mice (Figure 1D+E). ILC2 exhibited only a moderate reduction in the ability to produce IL-5 and IL-13 in response to infection (Figure 1F), however when coupled with decreased cellularity this led to an overall reduction in the number of cytokine producing ILC2 (Figure 1G). Thus, these data indicated that altering the availability of amino acids may modulate the quality and magnitude of the ILC2 response.

### ILC2 are poised for amino acid uptake

Our data suggested the induction of ILC2 responses may be sensitive to changes in the abundance of essential amino acids derived from dietary intake. Intriguingly, we observed that sort-purified ILC2 isolated from IL-33 treated mice exhibited a relative enrichment within their intracellular contents for many of the same amino acids (Figure 2A), including alanine, valine, leucine and isoleucine. To determine the underpinning molecular machinery through which alterations in amino acid abundance could potentially alter the ILC2 response, we examined published bulk RNA seq data (*22*) to analyse the expression of a range of solute carrier genes known to act as surface amino acid transporters in ILC2, in comparison to CCR6^+^ ILC3 (ILC3) (Figure 2B). We observed that ILC2, but not ILC3, constitutively expressed high levels of multiple solute carriers, most notably *Slc3a2, Slc7a5* and *Slc7a8*, known to encode for amino acid transporters (Figure 2B). *Slc3a2* encodes the protein CD98 – a chaperone molecule and heavy chain subunit that heterodimerises with other solute carriers to form active amino acid transporters, and ILC2 were also enriched for the CD98 binding partners *Slc7a5* and *Slc7a8*, which together form the surface large neutral amino acid transporters LAT1 and LAT2 respectively. In contrast, expression of other CD98 binding partners such as *Slc7a6, Slc7a7, Slc43a1* and *Slc43a2* were either not enriched in ILC2, or not detected in the data set (Figure 2B, Immgen database, (*22*)). LAT1 and LAT2 primarily transport a range of amino acid substrates with overlapping specificity to those also observed to be enriched in both the feces of helminth infected mice and enriched within the intracellular content of ILC2 (Figure1A+B, Figure 2A) (*23-26*), suggestive of a possible role for these transporters in ILC2 responses.

**Figure 2.**
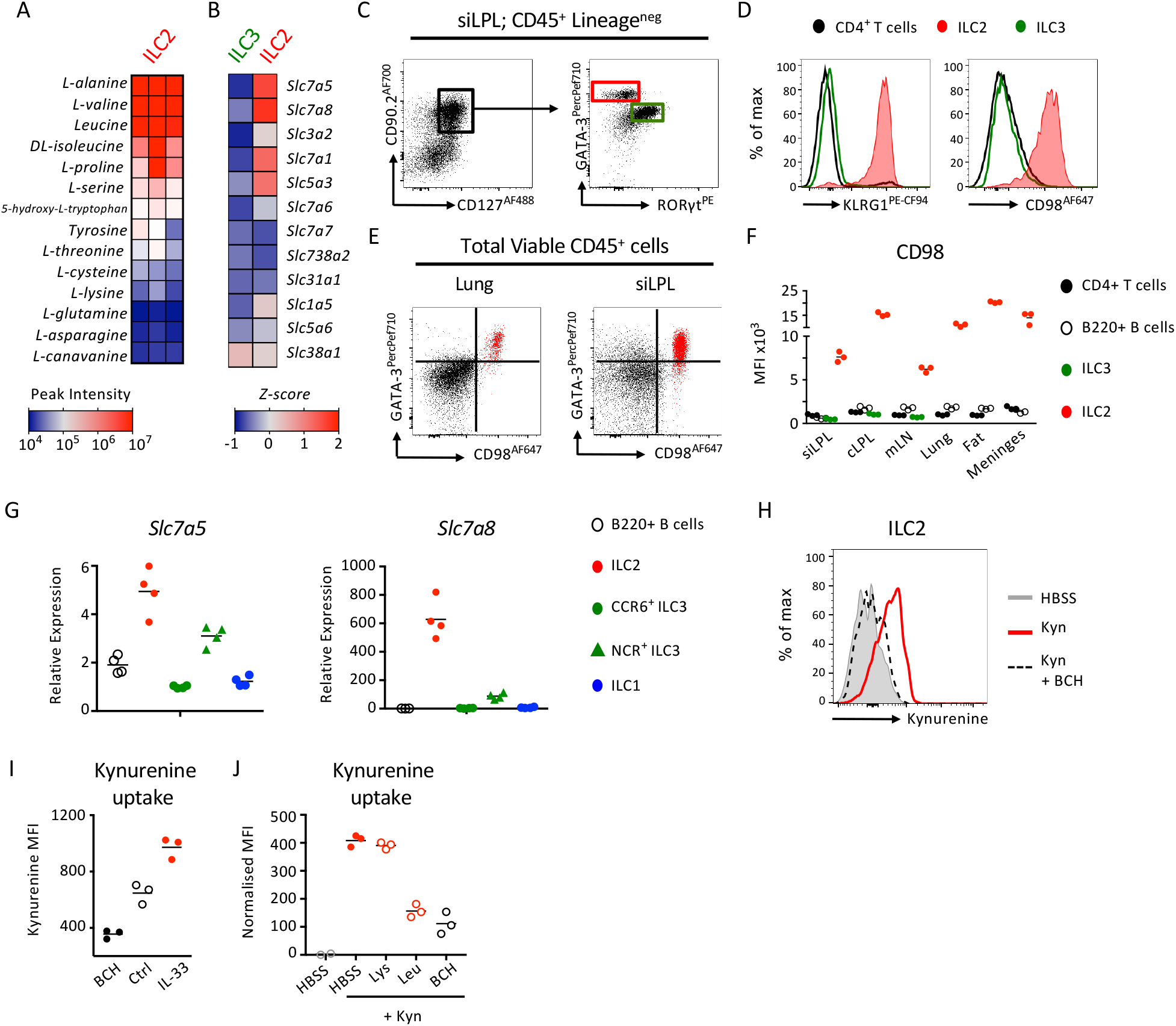
ILC2 are preferentially poised to import large neutral amino acids. A) Analysis of the intracellular amino acid content of ILC2 sort-purified from IL-33 treated mice (*n=*3 independent replicates of cells pooled from 2 mice and representative of two independent experiments). B) Comparison of mean expression of amino acid transporter-associated genes in ILC2 and CCR6^+^ ILC3 (ILC3) from public data (www.Immgen.org; (*22*)). C) Representative gating and D) surface expression of KLRG1 and CD98 on CD4^+^ T cells (black), ILC2 (red) and ILC3 (green) from small intestinal lamina propria (siLPL). E) Representative flow plots demonstrating co-expression of GATA-3 and CD98 in lung and siLPL amongst total CD45+ cells. F) Expression of CD98 on CD4^+^ T cells (black), B220^+^ B cells (white), ILC3 (green) and ILC2 (red) in siLPL, colon lamina propria (cLPL), mesenteric lymph node (mLN), lung, white adipose tissue (fat) and meninges. G) Relative expression of *Slc7a5* and *Slc7a8* in ILC subsets and B cells, normalised to CCR6^+^ ILC3 (*n=4* per group and representative of at least two independent experiments). H) Representative histogram of Kynurenine uptake in lung ILC2 incubated for 5 minutes with either HBSS alone (negative control), 200µM Kynurenine (Kyn) or Kynurenine plus 10mM BCH. I) Kynurenine uptake in ILC2 from naïve (Ctrl) or IL-33 treated mice, or incubated with Kynurenine and BCH (*n=3* per group and representative of two independent experiments). J) Kynurenine uptake in lung ILC2 in the presence or absence of excess (5mM) Lysine (Lys), Leucine (Leu) or 10mM BCH (*n*=3 replicates per condition, and representative of at least three independent experiments).

Consistent with this, we could detect constitutively elevated steady-state expression of CD98 on the cell surface of ILC2 – but not ILC3, CD4^+^ T cells or B220^+^ B cells – in a wide range of tissues (Figure 2C-F). Strikingly, we found a combination of GATA-3 and CD98 alone was sufficient to identify ILC2 amongst total CD45+ cells without prior lineage exclusion or pre-gating on classical ILC-associated markers (CD127, CD90.2), further indicating the preferentially heightened expression of CD98 by ILC2 amongst mucosal-resident lymphocytes (Figure 2E, Fig. S2A+B), while CD98 could also be detected on bone marrow ILC2 precursors (ILC2P; Fig. S2C). To validate whether surface CD98 expression on ILC2 was indicative of LAT activity we confirmed elevated expression of both *Slc7a5* and *Slc7a8* by RT-PCR in sort-purified ILC2 (Figure 2G). We then utilized a previously reported assay which utilizes the autofluorescent properties of the tryptophan metabolite kynurenine as a proxy of LAT transporter activity and amino acid uptake (*27*). Uptake of kynurenine was detected in naïve ILC2, which was inhibited by co-culture with the LAT-inhibitor BCH (Figure 2H) and found to be enhanced in ILC2 from IL-33 treated mice (Figure 2I). Kynurenine uptake in naïve ILC2 contrasted with CD4+ T cells which required TCR engagement to both upregulate surface CD98 and actively take up kynurenine (Fig. S2D, in line with previous findings (*27, 28*). Moreover, uptake of kynurenine by ILC2 was reduced by competition with excess levels of the high affinity LAT-substrates leucine and methionine, as well as alanine (a high affinity substrate of Slc7a8) but not lysine - which is not transported by Slc7a5 or Slc7a8 but rather by related y+LAT family members (Figure 2J and Fig. S2E) *(29-31)*. Together these findings indicate that ILC2 are preferentially poised to import amino acids via the surface large neutral amino acid transporters *Slc7a5* (LAT1) and *Slc7a8* (LAT2).

### Cell-intrinsic deletion of Slc7a5 or Slc7a8 impairs ILC2 expansion

As ILC2 were found to preferentially express CD98 along with two distinct partner chains *Slc7a5* (LAT1) and *Slc7a8* (LAT2), we next aimed to determine the role of these transporters during an ILC2 response. First, we generated mice with a conditional deletion of *Slc7a5* in ILC2 by crossing Red5^Cre^ mice (*32*) with *Slc7a5* ^fl/fl^ mice (*28*) (Figure 3A), and determined the effect on ILC2 responses following activation. ILC2 from IL-33 treated Red5^Slc7a5 fl/fl^ mice exhibited comparable expression of ST2 and KLRG1 as compared to Red5^Cre^ control animals but had a clear reduction in surface CD98 expression (Figure 3B). Moreover, while ILC2 frequencies at steady state were comparable between Red5^Cre^ and Red5^Slc7a5 fl/fl^ mice, ILC2 lacking cell-intrinsic *Slc7a5* expression demonstrated a clear defect in expansion following *in vivo* activation with IL-33 (Figure 3C+D). This correlated with a reduced percentage of cells expressing Ki-67 and notably, consistently reduced intensity of Ki-67 staining amongst positive cells (Figure 3E-G). In contrast, ILC2 exhibited comparable frequencies of IL-5 and IL- 13 positive cells in the absence of *Slc7a5* (Figure 3H+I), suggesting disruption of ILC2-intrinsic amino acid transport may largely perturb the magnitude but not the quality of the ILC2 response following activation.

**Figure 3.**
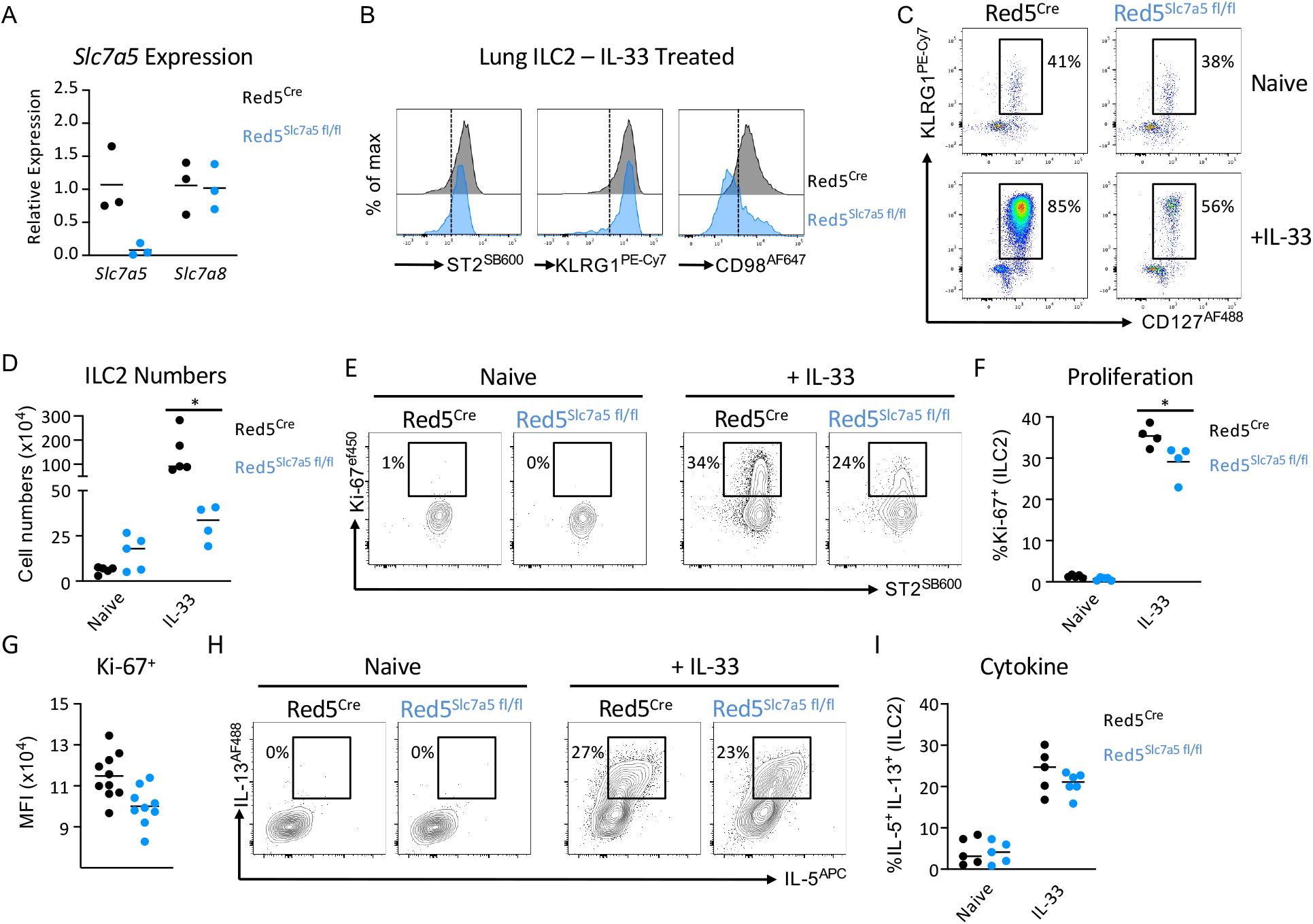
*Slc7a5* / LAT1 regulates the magnitude of ILC2 expansion following activation. A) Validation of *Slc7a5* deletion in Red5^Cre^ x *Slc7a5*^fl/fl^ mice (*n=3* technical replicates of sort-purified ILC2 pooled from the lungs of 2-3 IL-33 treated mice per replicate, representative of two independent experiments). B) Representative histograms of ST2, KLRG1 and CD98 in Red5^Cre^ control and Red5^Cre^ x *Slc7a5*^fl/fl^ mice (representative of 3-4 mice per group and at least three independent experiments). C) Frequencies and D) numbers of KLRG1+ CD127+ ILC2 (pre-gated on CD45+ Lineage negative cells) in the lungs of naïve or IL-33 treated Red5^Cre^ control and Red5^Cre^ x *Slc7a5*^fl/fl^ mice. (C+D, *n*=4-5 mice per group, representative of at least three independent experiments). E) Representative flow plots, F) quantification and G) mean fluorescent intensity of Ki-67 expression in ILC2 from control and IL-33 treated Red5^Cre^ control and Red5^Cre^ x *Slc7a5*^fl/fl^ mice. (E+F, *n*=4 mice per group, representative of three independent experiments, G, *n=*9-10 per group and pooled from two independent experiments) H) Representative flow plots and I) quantification of IL-5 and IL-13 producing ILC2 from control and IL-33 treated Red5^Cre^ control and Red5^Cre^ x *Slc7a5*^fl/fl^ mice. (H+I, *n*=5-6 mice per group, representative of at two independent experiments). Data shown as individual values and mean +/- SEM, * p< 0.05.

In contrast to *Slc7a5*, which has previously been attributed roles in the activation of other lymphocyte populations (*28, 33*), a role for *Slc7a8* in immune cells has not previously been described. Of note however, *Slc7a8* was previously listed amongst the top signature-defining genes of intestinal ILC2 by RNA sequencing (*22*) and was confirmed by RT-PCR to be highly and uniquely expressed in ILC2 derived from multiple tissues (Fig. S3A), but not in resting B or T cells (Fig. S3A+B), suggesting ILC2 may utilize multiple amino acid transporters to ensure a sufficient supply of these metabolic substrates. To test the role of *Slc7a8* in ILC2 responses we obtained and validated a *Slc7a8* knockout allele (Fig. S3B), which was subsequently converted to a loxP-flanked conditional allele via use of a FlpO recombinase. The Slc7a8 ^fl/fl^ allele was further backcrossed with Red5^Cre^ mice to generate Red5^Slc7a8 fl/fl^ animals with an ILC2-intrinsic deletion of *Slc7a8* (Red5^Slc7a8 fl/fl^). In contrast to our observations with IL-33 activated ILC2 (Figure 3B), and suggestive of a complimentary nature of these two amino acid transporters, we failed to observe any reduction in surface CD98 on steady state ILC2 in the absence of *Slc7a5* alone, whereas deletion of *Slc7a8* led to a reduced expression of surface CD98 in naïve ILC2 (Figure 4A). However confirming our previous findings, upon IL-33 activation *in vivo* CD98 expression on ILC2 was markedly reduced by the absence of *Slc7a5*, whereas in the absence of *Slc7a8* CD98 expression was in part maintained on the surface of ILC2 (Figure 4A). To try and reconcile these findings we determined the relative expression of the two transporter genes in naïve and IL-33 treated ILC2 and found that indeed *Slc7a8* was dominant in naïve animals but that the ratio between the two CD98 partner chains became relatively equal after activation (Figure 4B), suggestive of different contributions of *Slc7a5* and *Slc7a8* to functional amino acid transporter heterodimers in activated and resting ILC2. Nonetheless ILC2 frequencies and numbers were found to be comparable in naïve Red5^Slc7a8 fl/fl^ and littermate control animals. Instead a reduced expansion of ILC2 in response to IL-33 was observed in the absence of *Slc7a8* that was associated with a reduced frequency of Ki-67-expressing cells, but no cell-intrinsic defect in cytokine production (Figure 4C-G), comparable to results obtained with *Slc7a5* deletion. Thus, together these findings suggest that ILC2 express distinct large amino acid transporters with differing expression patterns in resting and activated cells, both of which act to support optimal cell expansion upon activation.

**Figure 4.**
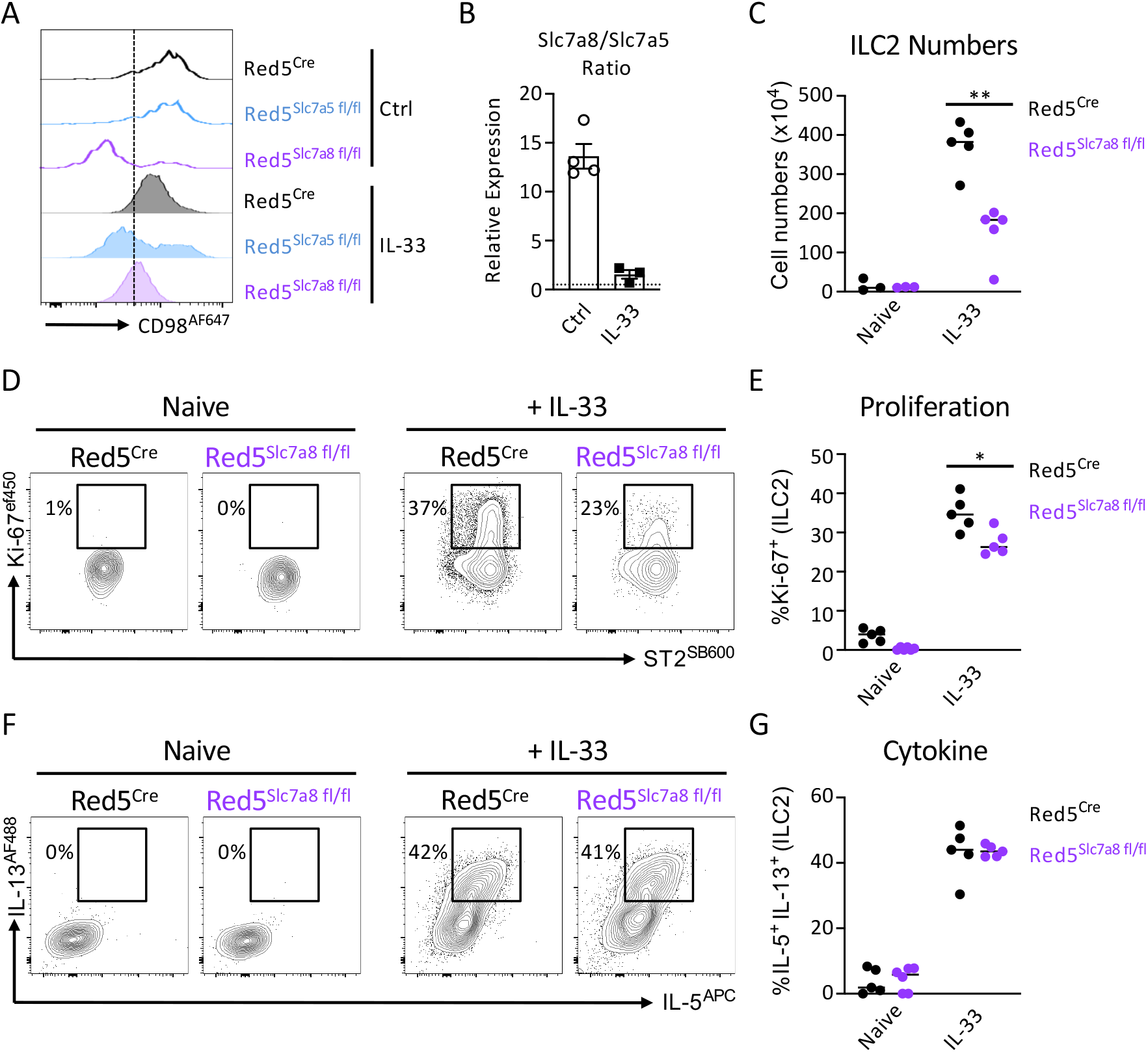
Differential expression of *Slc7a8* / LAT2 is required for optimal ILC2 expansion following activation. A) Representative histograms of CD98 expression in lung ILC2 from control (Ctrl) and IL-33 treated Red5^Cre^ control and Red5^Cre^ x *Slc7a8*^fl/fl^ mice (representative of *n*=3-5 mice per group and at least two independent experiments). B) Relative expression ratio of Slc7a8 to Slc7a5 in sort-purified ILC2 from control or IL-33 treated animals (*n=*3-4 technical replicates per group, representative of two independent experiments). C) ILC2 numbers, D) representative flow cytometry plots and E) quantification of Ki-67+ ILC2. F) Representative flow cytometry plots and G) quantification of IL-5+ IL-13+ ILC2 in lung ILC2 from control (Ctrl) and IL-33 treated Red5^Cre^ control and Red5^Cre^ x *Slc7a8*^fl/fl^ mice (*n*=3-5 mice per group and representative of at least two independent experiments). Data shown as individual values or mean +/- SEM, * p< 0.05, ** p< 0.01.

### Amino acid transporter deficiency impairs ILC2 responses to helminth infection

Our data demonstrate a reduced ability of ILC2 to expand and proliferate in response to IL-33 in the absence of either *Slc7a5* or *Slc7a8*. To determine the role of these transporters in generating ILC2 response to a more physiological infectious stimulus, we infected control or floxed mice with *N. brasiliensis* (Figure 5). In line with our prior findings, we noted that while naïve ILC2 had impaired surface CD98 expression in the absence of *Slc7a8* they increased compensatory solute carrier expression upon activation by helminth infection (Figure 5A). In contrast naïve ILC2 lacking *Slc7a5* exhibited comparable CD98 surface expression, but were unable to maintain surface CD98 upon activation by infection – again suggesting *Slc7a5* is proportionally increased and constitutes an elevated proportion of LAT heterodimers following ILC2 activation (Figure 5A). As with IL-33 activation, ILC2 expansion was reduced in the absence of either amino acid transporter following helminth infection (Figure 5B). Moreover, this reduced ILC2 response correlated with altered kinetics of infection. In particular, elevated worm burdens were observed in the intestines of mice at day 4 post infection as compared to control animals (Figure 5C+D). Notably, only mice lacking ILC2-intrinsic *Slc7a8* showed alterations in lung worm burdens at day 2, potentially suggesting a dominant role for this transporter in the early phase of an ILC2 response. Thus, in the context of a mucosal helminth infection the expression of the amino acid transporters LAT1 and/or LAT2 is required for optimal ILC2 responses.

**Figure 5.**
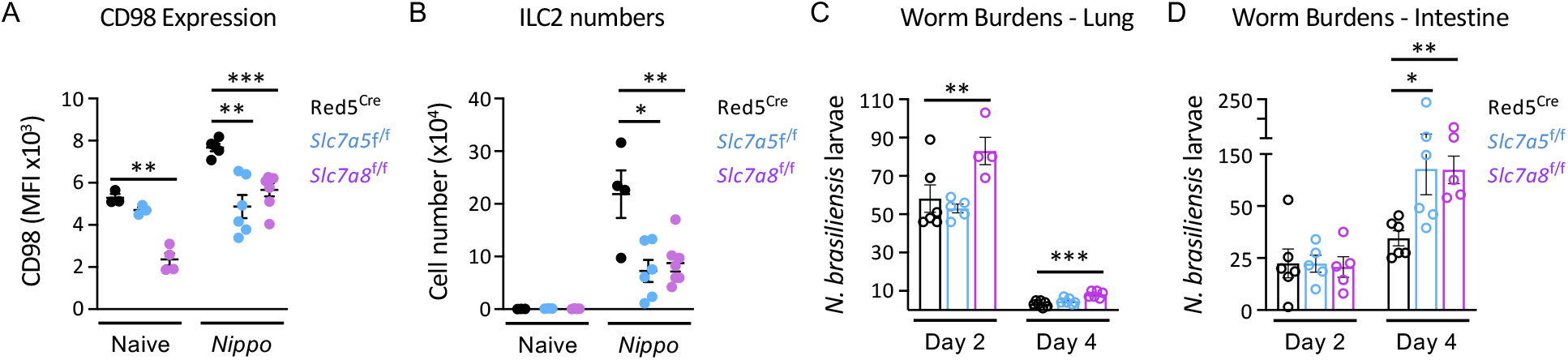
Disruption of large neutral amino acid transport in ILC2 dampens protection to *N. brasiliensis* infection. A) Mean Fluorescent Intensity of CD98 expression in lung ILC2, B) ILC2 cell numbers from control naïve and *N. brasiliensis* infected (day 7 p.i.) Red5^Cre^ x *Slc7a5*^fl/fl^ and Red5^Cre^ x *Slc7a8*^fl/fl^ mice (*n*=3-6 mice per group and representative of two independent experiments). Worm burdens in the C) lung and D) small intestine of *N. brasiliensis* infected (day 2+4 p.i.) Red5^Cre^ controls, Red5^Cre^ x *Slc7a5*^fl/fl^ and Red5^Cre^ x *Slc7a8*^fl/fl^ mice (*n*=4-6 mice per group and representative of two independent experiments). Data shown as individual values or mean +/- SEM, * p< 0.05, ** p< 0.01, *** p< 0.001.

### Perturbation of ILC2 amino acid transport results in metabolic stress

Ensuring a sufficient intracellular supply of amino acids is critical for cellular function, not only by providing the building blocks for the generation of biomass, but also via effects on cellular metabolism. Thus, we hypothesised that the consequences of perturbed amino acid transport would most likely be evident at the level of the proteome. To our knowledge proteomic analysis has not previously been attempted on ILC populations, therefore as a proof of concept we first sort-purified wild type ILC2 from IL-33 treated animals to determine feasibility. Using this approach, we were able to reproducibly detect over 5000 individual proteins from ILC2. Comparison of protein copy number with bulk RNA seq data of mRNA transcripts revealed a largely linear correlation between genes and their products, including classical ILC2 genes and proteins (Fig. S4A). However, in some cases (e.g. Thy1) the protein copy number diverged significantly from the relative gene expression level, indicating possible differences between transcriptomic and proteomic data in predicting ILC2 biology (Fig. S4A). Proteins associated with ILC2 phenotype and function, or cellular metabolism, were robustly detected but varied in their total copy number distribution (Fig. S4B). While these data demonstrate the feasibility of proteomic analysis of *in vivo* expanded ILC2, we were unable to generate sufficient material from naïve animals for comparison. Next, to investigate the role of amino acid transporter deletion on the ILC2 proteome we similarly sort-purified ILC2 from IL-33 treated Red5^Cre^, Red5^Slc7a5 fl/fl^ and Red5^Slc7a8 fl/fl^ mice and identified over 6000 proteins, of which ∼200 proteins differed significantly by genotype (Figure 6A). We confirmed efficient deletion of Slc7a5 and Slc7a8 protein in the respective knockout animals (Figure 6B), while ILC2 expression of activating cytokine receptors (Fig. S4C), transcription factors and canonical surface markers (Fig. S4D) were unchanged by transporter deletion. Via GO Term enrichment of the differentially expressed protein list we identified cell cycle progression, metabolism and protein translation as the major pathways perturbed in the absence of either amino acid transporter (Figure 6C).

**Figure 6.**
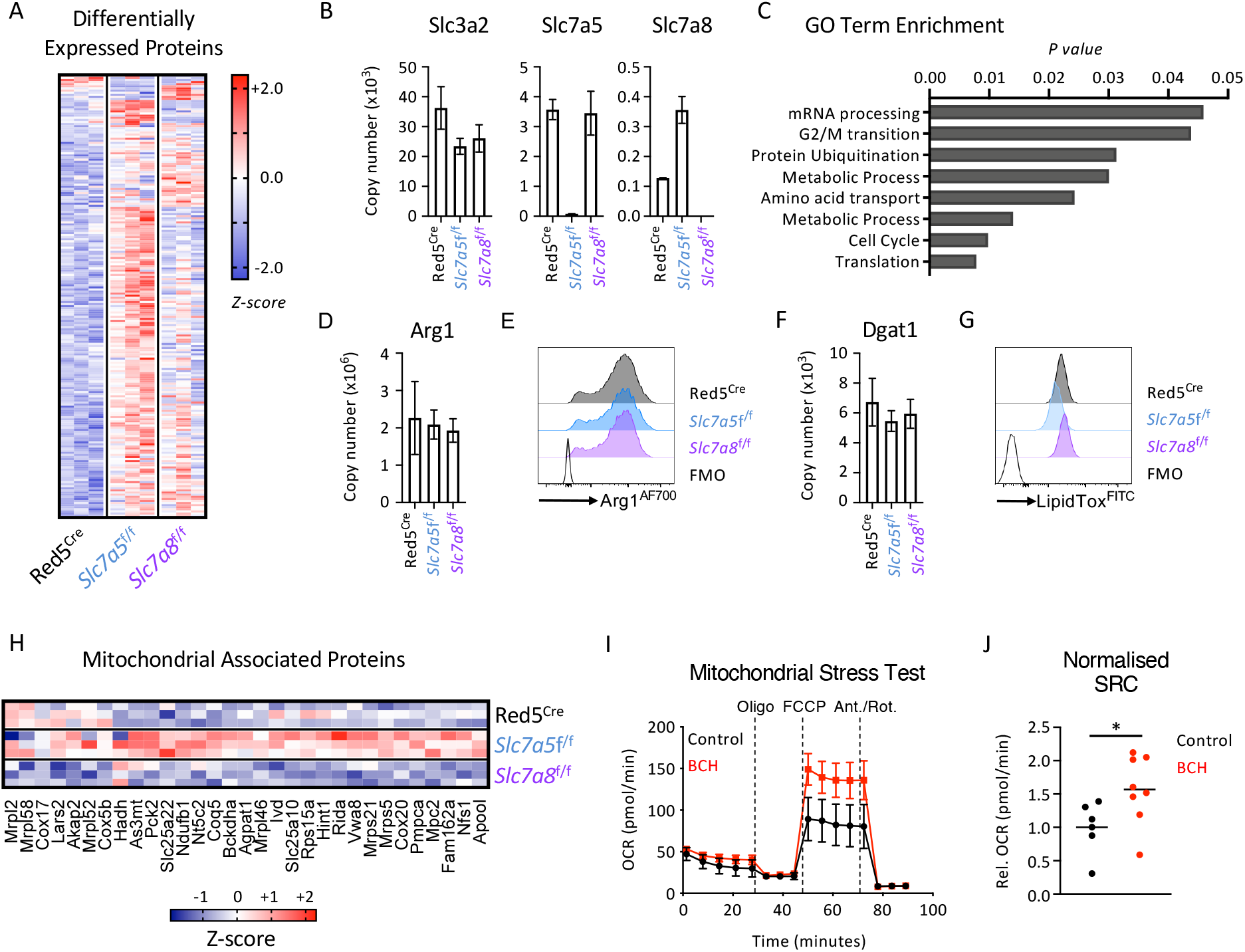
Proteomics of LAT-deficient ILC2 reveals metabolic imbalance. A) Differentially expressed proteins and B) copy numbers of target amino acid transporter associated proteins in sort-purified lung ILC2 from IL-33 treated Red5^Cre^ controls, Red5 x *Slc7a5*^fl/fl^ or Red5 x *Slc7a8*^fl/fl^ mice (*n*=3 replicates of cells pooled from 2-3 mice and representative of a single experiment). C) Go-term enrichment analysis of differentially expressed proteins across all genotypes in A. D) Arg1 protein copy number and E) flow cytometry analysis in lung ILC2 of IL-33 treated mice. F) Dgat1 copy number and G) intracellular lipid content (lipidTox staining) analysed by flow cytometry analysis in lung ILC2 of IL-33 treated mice (D+F, *n*=3 replicates of cells pooled from 2-3 mice and representative of a single experiment, E+G representative of at least *n=3* per genotype). H) Enrichment of mitochondrial associated proteins amongst differentially expressed proteins (identified with MitoCarta and MitoMiner). I) Extracellular flux analysis and J) spare respiratory capacity (SRC) of sort-purified ILC2 from IL-33 treated mice cultured with or without 10mM BCH overnight (*n*=3-4 technical replicates per experiment, I represents a single experiment, representative of two independent experiments, J representative of data pooled from two independent experiments). Data shown as individual values or mean +/- SEM, * p< 0.05.

Efficient nutrient uptake by ILC2 has previously been shown to act as a key determinant of cellular metabolism and the magnitude of the effector function, thus we investigated previously reported metabolic pathways implicated in the ILC2 response. However we found that Arginase-1 (Arg1) protein expression was not altered in the absence of amino acid transporter expression (Figure 6D+E)(*20*), nor was expression of Diacylglycerol acyltransferase 1 (Dgat1) (Figure 6F) or overall intracellular lipid storage (Figure 6G)(*18*). In contrast we identified protein signatures indicative of altered mitochondrial biology – especially in the absence of Slc7a5 – suggesting that lack of amino acid uptake may alter mitochondrial function of activated ILC2s (Fig. 6H). To test this, we sort-purified wild type ILC2 from IL-33 treated animals and cultured them overnight with the LAT-inhibitor BCH to impede LAT-dependent uptake of amino acids, and subsequently assessed the consequences via a Mitochondrial Stress Test. We consistently observed that ILC2 incubated with BCH exhibited a higher oxygen consumption rate upon addition of FCCP, which disrupts mitochondrial proton transport and ATP synthesis (Figure 6I), and had an increased spare respiratory capacity compared to control cells (Figure 6J), together suggesting that amino acid transporter blockade may lead to altered mitochondrial function, possibly as a compensatory measure in the context of perturbed intracellular amino acid availability.

### Intracellular amino acid availability controls proliferation via mTOR and metabolic rewiring

LAT-dependent intracellular amino acid availability has been extensively demonstrated to be a key regulator of activation of the mammalian target of rapamycin (mTOR), which in turn acts as a critical cellular hub that integrates nutrient availability with activating signals from growth factors, cytokines and other activating cues to determine downstream changes in cellular metabolism, protein translation, biomass synthesis and proliferation (23, 34, 35). To first determine the cues that activate mTOR in ILC2 under normal culture conditions we cultured sort-purified cells with cytokines, alarmins and neuropeptides known to influence ILC2 responses. As expected, we found that IL-7 poorly induced mTOR activation, as indicated by phosphorylation of ribosomal protein S6 (pS6), which is consistent with its role in homeostatic maintenance of ILCs (Figure 7A). In contrast, ILC2 cultured with IL-2, IL-25, IL-33 and Neuromedin U (NmU) all drove robust phosphorylation of S6, which could be completely or partially prevented by co-incubation with the mTOR inhibitor PP242 (Figure 7A+B). We then determined whether ablation of amino acid uptake with the LAT-inhibitor BCH could alter mTOR activation in response to an activating signal (IL-33), and indeed found that pre-incubation of ILC2 with BCH reduced pS6 in the presence of IL-33 in comparison to cells activated with IL-33 alone, although pS6 was only partially suppressed when compared to complete mTOR inhibition with PP242 (Figure 7C). Similarly, ILC2 cultured in leucine free media exhibited lower pS6 expression in response to IL-33 when compared to cells cultured in leucine replete media (Figure 7D). Together these findings suggest amino acid uptake via LATs on ILC2 acts in part to tune mTOR activation.

**Figure 7.**
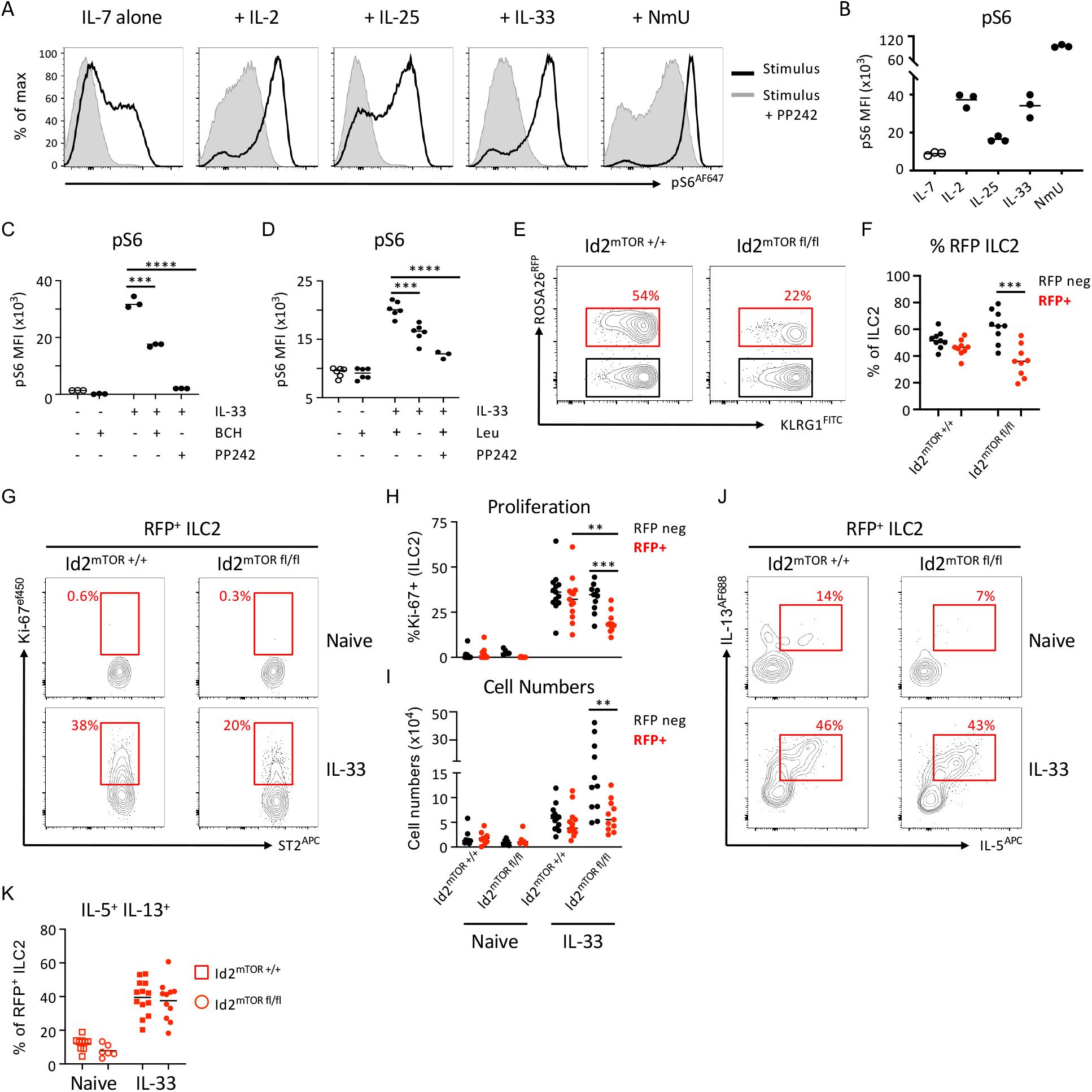
Regulation of mTOR activation by amino acid transport controls magnitude of ILC2 response. A) Representative flow plots and B) quantification of pS6 in sort-purified ILC2 cultured for 30 minutes in the presence of 20ng/ml IL-7 alone or IL-7 in combination with 20ng/ml IL-2, IL-25, IL-33 or 1µg/ml NmU, with or without the mTOR inhibitor PP242 (500nm). (*n*=3 technical replicates per condition, representative of two independent experiments). C+D) Phosphorylation of S6 in sort-purified ILC2 cultured with either C) IL-33 with or without a two hour pre-incubation with 10mM BCH (*n*=3 technical replicates per condition, representative of two independent experiments), or D) IL-33 in ILC2 cultured with either Leucine sufficient or deficient media (*n*=3-6 technical replicates per condition, representative of two independent experiments). E) Representative flow plots and F) quantification of RFP expression in IL-33 elicited lung ILC2 from Id2^ERT2Cre^ x Rosa26^tdRFP^ controls (Id2^mTOR+/+^) or Id2^ERT2Cre^ x Rosa26^tdRFP^ x mTOR^fl/fl^ mice (Id2^mTORfl/fl^) (*n*=9 per group, pooled from two independent experiments). G) Representative flow plots and H) quantification of Ki-67 expression and I) cell numbers of RFP negative and RFP+ lung ILC2 from naïve and IL-33 treated Id2^mTOR+/+^ and Id2^mTORfl/fl^ mice (*n*=5-9 per group for naïve mice and *n*=11-13 for IL-33 treated mice, pooled from three independent experiments). J) Representative flow plots and K) quantification of IL-5 and IL-13 expression in RFP+ ILC2 from naïve and IL-33 treated Id2^mTOR+/+^ and Id2^mTORfl/fl^ mice (*n*=5-9 per group for naïve mice and *n*=11-13 for IL-33 treated mice, pooled from three independent experiments). Data shown as individual values or mean +/- SEM, * p< 0.05, ** p< 0.01, *** p< 0.001, **** p< 0.0001.

Finally, as *Slc7a5* and *Slc7a8* primarily led to a reduced expansion of ILC2 without altering cell-intrinsic cytokine production, we determined to what extent mTOR regulation could contribute to these phenotypes. To circumnavigate potential developmental defects caused by a constitutive deletion of such a key sensing hub, we generated an inducible ERT2 Cre-driven model of mTOR deletion under the control of the *Id2* locus (to predominantly target ILCs), which upon tamoxifen administration drove both an RFP reporter allele as previously described (*36, 37*), and deletion of flanking loxP sites in mTOR (Id2^mTOR fl/fl^). Following tamoxifen administration and activation of ILC2 via IL-33 we noted a reduced frequency of RFP expressing cells in Id2^mTOR fl/fl^ mice when compared to Id2^mTOR +/+^ control mice, indicating Cre-activated cells (RFP^+^) may be at a competitive disadvantage to wild type (RFP^-^) cells in the absence of mTOR (Figure 7E+F). In line with this, we observed an intrinsic defect in Ki-67 expression and proliferation amongst Cre-activated RFP^+^ cells from Id2^mTOR fl/fl^ when compared to otherwise mTOR competent RFP^-^ cells in which Cre had not been recombined in the same mice (Figure 7G-I). In contrast, proliferation and cell numbers were not perturbed by Cre activation and RFP expression in mice lacking the floxed allele (Figure 7G-I). Mirroring our findings with amino acid transporter deletion in ILC2, RFP^+^ cells lacking mTOR showed no defect in cytokine production following IL-33 activation when compared to those in control mice (Figure 7J-K). Thus, our findings suggest amino acid uptake via LATs mat regulate the magnitude of the ILC2 expansion via tuning of mTOR activation upon activation.

Collectively, the findings presented here suggest that ILC2 - unlike other major lymphocyte populations such as T cells - are pre-poised for the uptake of amino acids from the tissue environment in order to fuel optimal proliferation and cell expansion upon activation. This poised state appears to be in part due to the preferential expression of *Slc7a8* in ILC2 at steady state, which was otherwise not detected in resting T cells. This suggests ILC2 may employ two distinct large neutral amino acid transporters, both prior to and following activation, to ensure sufficient intracellular amino acid availability to fuel a rapid innate response. One possibility is that the differing substrate specificity of the two transporters facilitates differential uptake of amino acids. Indeed, *Slc7a8* has been suggested to have higher specificity for alanine (*31*), an amino acid found to be the most highly enriched in sort-purified ILC2 (Figure 2A), and which has recently also been shown to regulate mTOR activation in addition to classical substrates such as leucine (*38*). To our knowledge this is the first report of a role for *Slc7a8*/LAT2 in immune cell functionality.

In contrast, we noted a relative increase of *Slc7a5* expression and associated dependence on *Slc7a5* expression for surface CD98 following activation, similar to that reported for activated T cells (*28*). While this data suggests *Slc7a8*/LAT2 may preferentially act in a steady state setting we found no differences in ILC2 frequency or numbers across tissues in naïve animals, although ILC2 lacking *Slc7a8* failed to proliferate and expand in response to IL-33. One possibility is that *Slc7a8* may determine metabolic tone or innate fitness of naïve ILC2 to prime them for rapid proliferation, although due to the limitations in performing extensive molecular and cellular analysis of naïve ILC2 we have been unable to investigate the different contributions of LAT1 and LAT2 in resting ILC2 further within the scope of this study. Nonetheless, and in line with our findings, a recent report similarly demonstrated that human ILC2 isolated from peripheral blood are also uniquely poised for amino acid uptake and that ILC2 cultured with inhibitors of downstream pathways associated with amino acid metabolism exhibited reduced cellular fitness and proliferation (*39*). This highlights a conserved requirement for amino acid uptake in ILC2 across species. Further studies and refined methodologies are needed to definitively dissect the different contributions of *Slc7a5* and *Slc7a8* to rare immune cell population biology and metabolism.

Finally, we observed that helminth infections increase the abundance of essential amino acids within the feces, in line with a previous report (*40*), potentially linking changes in environmental cues with the metabolic and proliferative capacity of the responding innate immune cell. It is tempting to speculate that ILC2 may express multiple amino acid transporters not only to ensure sufficient import of metabolic substrates required to underpin their rapid innate expansion and functionality, but also to enhance their sensitivity to environmental changes associated with infections that likely acted as a key evolutionary pressure to drive the emergence of this arm of the immune system. It is increasingly clear that a broad range of microbial and dietary metabolites regulate the activation of ILC2 (*17-20, 39*), and together with classical activating signals, such as alarmins and neuropeptides, nutrient and metabolite availability likely act as a further regulatory layer to tune the magnitude of the immune response within the tissue microenvironment and facilitate rapid innate immune responses.

## Materials and Methods

### Mice

Six to eight week old female C57BL/6 were purchased from Envigo, Cambridge, UK. Red5^Cre^ (B6(C)-*Il5*^tm1.1(iCre)Lky/^J, stock number 030926, originally generated by Richard Locksley, UCSF), Id2ERT2^Cre^ (B6.129S(Cg)-*Id2*^tm1.1(*Cre*/ERT2)Blh^/ZhuJ, stock number 016222, originally generated by Yuan Zhuang, Duke University) and mTOR^fl/fl^ mice (B6.129S4-*Mtor*^tm1.2Koz^/J, stock number 011009, originally generated by Sara Kozma, University of Cincinatti) were originally imported from Jackson laboratories. ROSA26^tdRFP^ were originally a kind gift from Hans Joerg Fehling, *Slc7a5*^fl/fl^ mice (B6.129P2-*Slc7a5*^tm1.1Daca^/J) were a kind gift from Doreen Cantrell (University of Dundee). *Slc7a8*^fl/fl^ mice were generated by crossing C57BL/6N-*Slc7a8*^tm2a(EUCOMM)*Hmgu*^/BayMmucd mice with mice containing a FlpO recombinase allele to remove the *lacZ* and Neomycin cassettes (KOMP/MMRRC repository, stock number 041243-UCD originally generated by Arthur Beaudet, Baylor College of Medicine), generating flanking loxP sites spanning the critical exon. In some experiments mice were fed a diet containing 21% protein or 5% protein for two weeks prior to infection or subsequent manipulation, diets were purchased from Envigo laboratories (TD. 140918 and TD. 140711). For activation of inducible Cre alleles mice were orally gavaged 5mg Tamoxifen in Corn oil every 2-3 days for a period of two weeks and rested one week prior to further experimental manipulation. For transgenic animal studies age- and sex-matched littermate controls were used within experiments where possible, mice were maintained at University of Manchester under specific pathogen free conditions, with water and chow provided *ad* libitum, with constant temperature and 12 hour light and dark cycle. All experiments were performed under license of the U.K. Home Office and under approved protocols. All animal studies were ethically reviewed and carried out in accordance with Animals (Scientific Procedures) Act 1986 and the GSK Policy on the Care, Welfare and Treatment of Animals.

### In vivo IL-33 treatment

Mice were injected intraperitoneally with 0.5µg of recombinant IL-33 (BioTechne) on day 0, 2 and 4 unless otherwise indicated. To maximise cell yield for sort-purification of ILC2 and *ex vivo* assays mice received additional doses of IL-33 and/or a higher dosing regimen (1µg).

### Helminth infections

Mice were infected with 300 L3 *Nippostrongylus brasiliensis* via subcutaneous injection, or 250 infective larvae of either *Heligmosomoides polygyrus* or *Trichnella spiralis* via oral gavage. Helminth life cycles were maintained and infective larvae kindly provided by the groups of Judi Allen, Richard Grencis and John Grainger at the University of Manchester.

### Tissue processing

Briefly, Lungs were collected in 2ml of PBS and thoroughly minced prior to the addition of 2mg/ml Collagenase D and 33µg/ml DNase. Tissue was shaken at 37C for 40 minutes at 200rpm, prior to mechanical disruption and passing over a 70µm nylon filter and flushing of remaining tissue with PBS. Supernatants were pelleted and cells briefly incubated with 2ml ACK buffer to lyse residual red blood cells, prior to being washed and resuspended in PBS containing 5% FCS and 1mM EDTA for flow cytometry staining. Mesenteric lymph nodes (mLN) were processed in a similar manner without enzymatic digestion, via manual disruption over a 70µm nylon filter. Intestinal lamina propria preparations were isolated by removing all fat and Peyer’s patch from intestines, opening longitudinally and flushing in PBS, followed by extensive vigorous vortexing of intestinal tissue in PBS and subsequent rounds of incubation and constant shaking with PBS containing 5% FCS, 1mM EDTA and 1mM DTT at 37C to remove mucus and epithelium. The remaining tissue was then incubated with constant shaking at 37C in RPMI media containing 0.1mg/ml collagenase/dispase (Roche) and 20µg/ml DNase (Sigma-Aldrich) for 45 minutes. Supernatant containing liberated lymphocytes was collected by passing tissue over a 70µm nylon filter, and cells pelleted and resuspended in PBS containing 5% FCS and 1mM EDTA for flow cytometry staining.

### Flow cytometry and cell sorting

Surface antibody staining was performed in PBS containing 5% FCS and 1mM EDTA and using a Fixable Aqua Dead Cell (Invitrogen) to determine viability. Cells were stained with the following cell surface antibodies and using the conjugates indicated in the figure labels and utilized for analysis with a BD Fortessa or cell-sorting with a BD Aria Influx; CD127 (IL-7Rα, Brilliant Violet 421, PE, or FITC, eBioscience; clone A7R34), ST2 (IL-33R Biotin; eBioscience, clone RMST2-33), CD45 (brilliant violet 650; clone 30-F11, BioLegend), CD3 (PerCP-Cyanine 5.5 or PE/Cy7; clone 145-2C11), CD5 (PerCP-Cyanine 5.5 or PE/Cy7, BioLegend; clone 53-7.3) NK1.1 (PerPC-Cyanine 5.5, BV395, or PE/Cy7, eBioscience; clone PK136), CD90-2 (alexa fluor 700 AM; clone 30-H12, BioLegend), B220 (CD45R, APC-e Fluor 780, eBioscience; clone RA3-6B2) CD11b (super bright 600, Invitrogen, APC-e Fluor 780; clone M1/70) CD11c (APC-e Fluor 780, eBioscience; clone N418), CD4 (super bright 600, eBioscience; clone RM4-5; BV395, eBioscience; clone GK1.5), SA-APC (streptavidin APC, eBioscience), SA-SB600 (streptavidin super bright 600, eBioscience), CD98 (Alexa Fluor 647; clone RL388, BioLegend), CD8α (FITC; clone 53-6.7, BioLegend), KLRG1 (PeCyanine 7, Invitrogen, FITC, or Pe-eFlour 610, eBioscience; clone 2F1), MHCII (eFluor 450; clone M5/114,15.2, Invitrogen).

Intracellular staining was performed by fixing cells for 30 minutes FoxP3 fix/perm buffers (eBioscience) prior to staining for 30 minutes in permeabilisation buffer (eBioscience) at 4C. Alternatively in order to retain reporter signals cells were first fixed with BD Cytofix/Cytoperm buffer (BD Biosciences) for 1 hour at 4C prior to staining intracellular antigens overnight at room temperature. Intracellular antbodies utilized in this study RORγt (PE; clone B2D), Ki-67 (eFluor 450, eBioscience; clone SolA1s), GATA 3 (PercP eFluor 710, eBioscience; clone TWAJ), Arg1 (Alexa fluor 700, eBioscience; clone A1exF5), IL-5 (PE or APC, eBioscience; clone TRFK5, BioLegend), and IL-13 (Alexa Fluor 488, or PeCyanine 7, Invitrogen; or eFluor 660, eBioscience; clone eBio13A). For phosphoFlow, stimulated cells were fixed with pre-warmed Phosflow Lyse/Fix buffer (BD Biosciences) for 10 minutes, washed and permeabilised with ice cold Perm Buffer III (BD biosciences) for 30 minutes, and subsequently stained with pS6 (Ser235 Ser236; APC, eBioscience; clone cupk43k). The Kynurenine uptake assay was performed as previously described (*27*).

### RT-PCR and Bulk RNA sequencing

Total RNA was purified using the RNeasy Micro Kit (Qiagen) and cDNA was prepared using the high capacity cDNA reverse transcription kit (Applied Biosystems). Real-time qPCR was performed with the real-time PCR StepOnePlus system (Applied Biosystems). Bulk RNA Seq of wild type ILC2, RNA was isolated from sort-purified cells, as above, and library preparation and bulk RNA sequencing was performed commercially with Novogene (UK) Company Ltd. Briefly, normalised RNA was used to generate libraries using NEB Next Ultra RNA library Prep Kit (Illumina). Indices were included to multiplex samples and mRNA was purified from total RNA using poly-T oligo-attached magnetic beads. After fragmentation, the first strand cDNA was synthesised using random hexamer primers followed by second strand cDNA synthesis. Following end repair, A-tailing, adaptor ligation and size section libraries were further amplified and purified and insert size validated on an Agilent 2100, and quantified using quantitative PCR (qPCR). Libraries were then sequenced on an Illumina NovaSeq 6000 S4 flowcell with PE150 according to results from library quality control and expected data volume.

### Extracellular flux analysis

ILC2 were sort-purified from IL-33 treated mice and incubated overnight with or without 10mM BCH. Seahorse plates and cartridges were prepared 18h before by adding 200µl XF Calibrant to each well (Seahorse Bioscience/Agilent, USA) to emerge probes, and incubating at 37C to calibrate. ILC2 were washed and plated onto poly-D-lysine-coated XF96 plates with XF RPMI media and rested for 30 minutes at 37C prior to analysis. For the mitochondrial stress test, Seahorse medium was supplemented with 25mM glucose (Thermo Scientific), 1mM sodium pyruvate and 2mM L-glutamine (Sigma Aldrich) and pH adjusted to 7.4. Cellular bioenergetics were assessed at 5-min intervals following sequential addition of 2µM Oligomycin, 2µM FCCP, 0.5µM Antimycin A and 0.5µM Rotenone (all Sigma-Aldrich) using an XF96e extracellular flux analyzer (Seahorse Bioscience/Agilent, USA) via sequential addition of 2µM Oligomycin, 1.5µM FCCP, 0.5µM Antimycin A and 0.5µM Rotenone (all Sigma Aldrich).

### Fecal metabolomics

The metabolic profiles of fecal samples were measured using ^1^H nuclear magnetic resonance (NMR) spectroscopy as previously described (*41*). Briefly, fecal samples (30 mg) were defrosted and combined with 600µL of water and zirconium beads (0.45 g). Samples were homogenized with a Precellys 24 instrument (45 s per cycle, speed 6500, 2 cycles) and spun at 14,000 *g* for 10 minutes. The supernatants (400µL) were combined with 250µL phosphate buffer (pH 7.4, 100% D_2_O containing 3 mM NaN_3_, and 1 mM of 3-(trimethyl-silyl)-[2,2,3,3-^2^H4]-propionic acid [TSP] for the chemical shift reference at δ0.0), before vortexing and centrifugation at 14,000 *g* for 10 minutes and transfer to 5 mm NMR tubes. All samples were anlaysed on a Bruker 700 MHz spectrometer equipped with a cryoprobe (Bruker Biospin, Karlsruhe, Germany) operating at 300 K. ^1^H NMR spectra were acquired for each sample using a standard one-dimensional pulse sequence using the first increment of the NOE pulse sequence for water suppression as previously described (*42*). Raw spectra were automatically phased, baseline corrected and calibrated to TSP using Topspin 3.2 (Bruker Biospin) and then digitized in a Matlab environment (Version 2018; Mathworks Inc, USA) using in-house scripts. Redundant spectral regions (related to water and TSP resonance) were removed, and the spectral data was manually aligned and normalized to the probabilistic quotient using in-house Matlab scripts. Peak integrals (relating to relative abundance) for metabolites of interest were calculated for each sample.

### Proteomics and Mass Spectrometry

For initial establishment of proteomic methodology, technical replicate pools of 5 million ILC2 were sort-purified from IL-33 treated mice. For comparison of control and conditional knockout mice, technical replicates averaging between 500,000 - 1 million ILC2 pooled from two individual mice were sort purified. Cell pellets were washed extensively with PBS to remove residual FCS and snap frozen. Samples were prepared for mass spectrometry by adding 100 µl of lysis buffer (5 % sodium dodecyl sulphate, 50 mM TEAB pH 8.5, 10 mM TCEP) to each cell pellet and shaking at 1000 rpm at room temperature for 5 minutes. Lysates were boiled for 5 minutes at 95 °C, sonicated for 15 cycles of 30 seconds each and treated with 1 µl benzonase for 15 minutes at 37 °C. Protein yield was determined using the EZQ protein quantitation kit (ThermoFisher Scientific) according to manufacturer’s instructions. Lysates were alkylated with 20 mM iodoacetamide for 1 hour at room temperature in the dark. Protein lysates were loaded on to S-Trap micro columns (ProtiFi) following the manufacturer’s instructions. Proteins were digested with 20:1 protein:trypsin (Trypsin Gold, Promega) in 50 mM ammonium bicarbonate for 3 hours at 47 °C before adding an additional 1 µg of trypsin and digesting for a further 1 hour at 47 °C. Peptides were eluted from columns and dried by SpeedVac and resuspended in 1 % formic acid at a peptide concentration of 0.1 µg/µl.

For LC-MS analysis of wild type ILC2 1.5 µg of peptide for each sample was analysed on a Q-Exactive-HF-X (Thermo Scientific) mass spectrometer coupled with a Dionex Ultimate 3000 RS (Thermo Scientific). The following LC buffers were used: buffer A (0.1% formic acid in Milli-Q water (v/v)) and buffer B (80% acetonitrile and 0.1% formic acid in Milli-Q water (v/v)). 1.5 μg aliquot of each sample was loaded at 15 μL/min onto a trap column (100 μm × 2 cm, PepMap nanoViper C18 column, 5 μm, 100 Å, Thermo Scientific) equilibrated in 0.1% trifluoroacetic acid (TFA). The trap column was washed for 3 min at the same flow rate with 0.1% TFA then switched in-line with a Thermo Scientific, resolving C18 column (75 μm × 50 cm, PepMap RSLC C18 column, 2 μm, 100 Å). Peptides were eluted from the column at a constant flow rate of 300 nl/min with a linear gradient from 3% buffer B to 6% buffer B in 5 min, then from 6% buffer B to 35% buffer B in 115 min, and finally to 80% buffer B within 7 min. The column was then washed with 80% buffer B for 4 min and re-equilibrated in 3% buffer B for 15 min. Two blanks were run between each sample to reduce carry-over. The column was kept at a constant temperature of 50°C at all times. Data was acquired using an easy spray source operated in positive mode with spray voltage at 1.9 kV, the capillary temperature at 250 °C and the funnel RF at 60 °C. The MS was operated in DIA mode using parameters previously described (*43*), with some modifications. A scan cycle comprised a full MS scan (m/z range from 350-1650, with a maximum ion injection time of 20 ms, a resolution of 120 000 and automatic gain control (AGC) value of 5 × 106). MS survey scan was followed by MS/MS DIA scan events using the following parameters: default charge state of 3, resolution 30.000, maximum ion injection time 55 ms, AGC 3 × 106, stepped normalized collision energy 25.5, 27 and 30, fixed first mass 200 m/z. Data for both MS and MS/MS scans were acquired in profile mode.

For conditional knockout LC-MS analysis, peptides were analysed on a Q Exactive™ plus, Mass Spectrometer (Thermo Scientific) coupled to a Dionex Ultimate 3000 RS (Thermo Scientific). The following LC buffers were used: buffer A (0.1 % formic acid in Milli-Q water (v/v)) and buffer B (80 % acetonitrile and 0.1 % formic acid in Milli-Q water (v/v). An equivalent of 1.5 µg of each sample was loaded at 10 μL/min onto a µPAC trapping C18 column (Pharmafluidics). The trapping column was washed for 6 min at the same flow rate with 0.1 % TFA and then switched in-line with a Pharma Fluidics, 200 cm, µPAC nanoLC C18 column. The column was equilibrated at a flow rate of 300 nl/min for 30 min. The peptides were eluted from the column at a constant flow rate of 300 nl/min with a linear gradient from 1 % buffer B to 3.8 % buffer B in 6 min, from 3.8 % B to 12.5 % buffer B in 40 min, from 12.5 % buffer B to 41.3 % buffer B within 176 min and then from 41.3 % buffer B to 61.3 % buffer B in 14 min. The gradient was finally increased from 61.3 % buffer B to 100 % buffer B in 1 min, and the column was then washed at 100 % buffer B for 10 min. Two blanks were run between each sample to reduce carry-over. The column was kept at a constant temperature of 50 °C.

Q-exactive plus was operated in positive ionization mode using an easy spray source. The source voltage was set to 2.2 Kv and the capillary temperature was 275 °C. Data were acquired in Data Independent Acquisition Mode as previously described (Doellinger et al., 2020), with some modifications. A scan cycle comprised of a full MS scan (m/z range from 345-1155), resolution was set to 70,000, AGC target 3 × 10^6^, maximum injection time 200 ms. MS survey scans were followed by DIA scans of dynamic window widths with an overlap of 0.5 Th. DIA spectra were recorded at a resolution of 17,500 at 200 m/z using an automatic gain control target of 3 × 10^6^, a maximum injection time of 55 ms and a first fixed mass of 200 m/z. Normalised collision energy was set to 25 % with a default charge state set at 3. Data for both MS scan and MS/MS DIA scan events were acquired in profile mode.

Raw mass spectrometry data was processed using Spectronaut (Biognosys; version 14.5.200813.47784 for wild type ILC2 and version 14.10.201222.47784 for conditional knockout comparisons). For all searches the DirectDIA option was selected. The following parameters were chosen: cleavage rules were set to Trypsin/P, maximum peptide length 52 amino acids, minimum peptide length 7 amino acids, maximum missed cleavages 2 and calibration mode automatic. Carbamidomethylation of cysteine was set as a fixed modification while the following variable modifications were selected: oxidation of methionine, deamidation of asparagine and glutamine and acetylation of the protein N-terminus. The FDR threshold for both precursor and protein was set at 1 %. DirectDIA data were searched against a mouse database from Uniprot release 2020 06. This database consisted of all manually annotated mouse SwissProt entries along with mouse TrEMBL entries with protein level evidence and a manually annotated homologue within the human SwissProt database. Estimates of protein copy number per cell were calculated using the histone ruler method (*44*).

### Statistics

Data presented as mean +/- SEM, unless indicated otherwise. Statistical analyses were performed using either Student’s t-test, Mann-Whitney test, Kruskal-Wallis test or one-way ANOVA, as indicated, and unless otherwise specified.

## Author contributions

MRH conceptualised, designed and performed experiments, wrote the manuscript and secured funding for the project. RKG contributed to conceptualisation and design and provided expert input. SH, MZK, IK and JK designed, performed and analysed experiments and associated data. LVS provided expert input and assisted with experimental design and provided mice, reagents and protocols. AJMH performed proteomic experiments and analysed data. GP and JRS performed fecal metabolomic experiments and analysis.

## Acknowledgements

The authors acknowledge members of the Hepworth lab for critical discussion, Gareth Howell, David Chapman and the University of Manchester flow cytometry core for support, the University of Manchester BSF staff for support with animal husbandry and maintenance, and David Knight, George Taylor and the University of Manchester Biological Mass Spectrometry Facility. We also thank the Fingerprints Proteomics Facility in the School of Life Sciences, University of Dundee for support. We also thank Naren Srinivasan and GlaxoSmithKline for financial and scientific support as part of an MRC CASE PhD studentship awarded to Suzanne Hodge. Research in the Hepworth Laboratory is supported by a Sir Henry Dale Fellowship jointly funded by the Wellcome Trust and the Royal Society (Grant Number 105644/Z/14/Z), a BBSRC responsive mode grant (BB/T014482/1) and a Lister Institute of Preventative Medicine Prize. Linda Sinclair was supported by a Wellcome Trust Principal Research Fellowship, awarded to Doreen Cantrell (Grant Number 205023/Z/16/Z).

