## Supplementary Figures for "Amino acid availability acts as a metabolic rheostat to determine the magnitude of ILC2 responses"

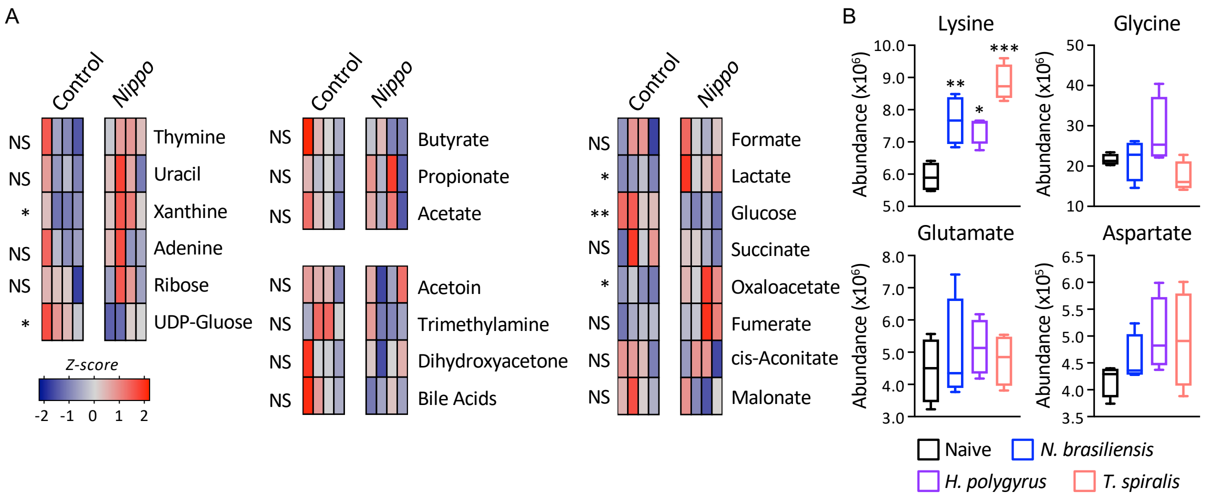


**Supplementary Figure 1. Changes in metabolite and dietary factors following helminth infection.** A) Relative abundance of fecal metabolites in control and day 7 post infection *N. brasiliensis* infected C57BL/6 mice (*n*=4 mice per group, representative of two independent experiments, data shows z-scores). B) Relative abundance of amino acids (related to Figure 1A) in naïve mice or mice infected with *N. brasiliensis* (day 7 p.i. blue), *H. polygyrus* (day 7 p.i., purple) or *T. spiralis* (day 7 p.i., pink), (*n*=4 mice per group, representative of one experiment, data shows relative abundance). Data shown as individual values or mean +/- SEM, * p< 0.05, ** p< 0.01, *** p< 0.001.

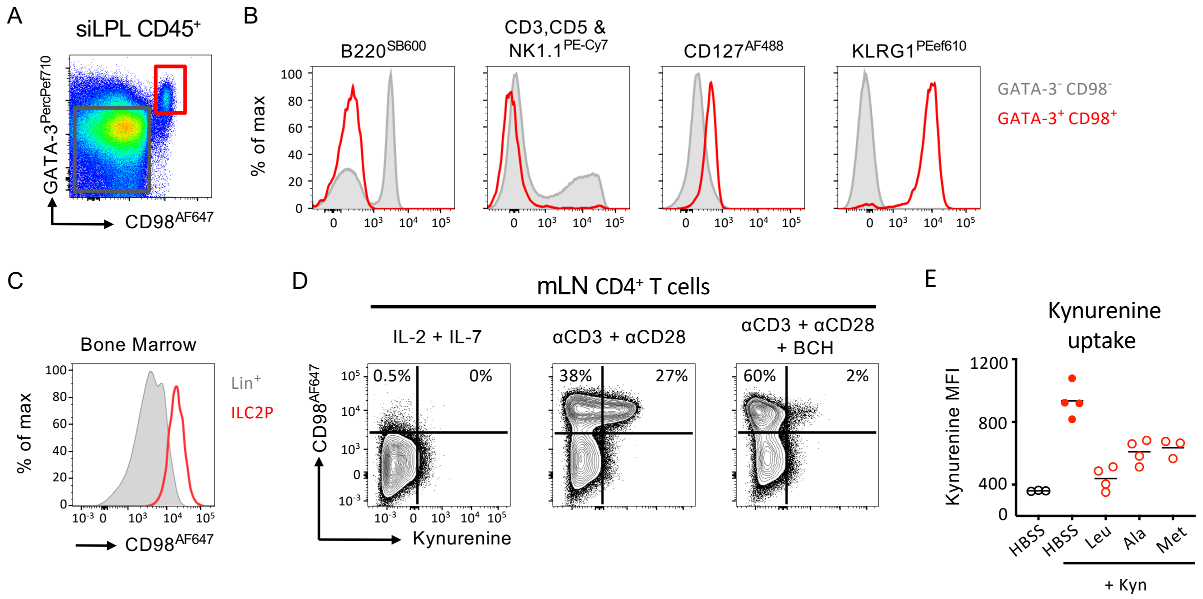


**Supplementary Figure 2. Amino acid transport in ILC2 and CD4+ T cells.** A+B) Enrichment of ILCs by CD98+ GATA-3+ gating from total CD45+ lymphocytes (representative of at least 3 independent experiments with 4-5 mice per group). C) CD98 expression on CD127+ CD25+ ST2+ ILC2 progenitors (ILC2P; red) and Lineage positive bone marrow cells (Lin$^+$ grey) (representative of *n*=4 mice per group and one experiment). D) Exemplar CD98 expression and Kynurenine uptake in CD4+ T cells from the mesenteric lymph node (mLN) cultured with either rIL-2 and rIL-7 alone, anti-CD3 and anti-CD28 monoclonal antibodies, or anti-CD3 and anti-CD28 plus BCH (representative of two independent experiments with *n*=2-4 replicate wells per group). E) Kynurenine uptake in lung ILC2 in the presence or absence of excess Leucine (Leu), Alanine (Ala) or Methionine (Met) (*n*= 3-4 replicates per condition, and representative of two independent experiments).

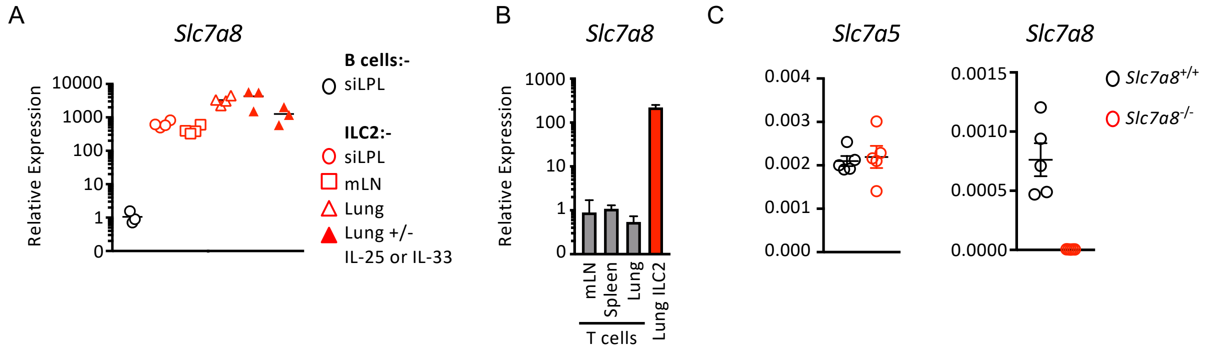


**Supplementary Figure 3. Selective *Slc7a8* expression in ILC2 and validation of deletion.** A) Relative expression of *Slc7a8* in sort purified ILC2 from the small intestinal lamina propria (siLPL), mesenteric lymph node (mLN), Lung, or Lung from mice previously treated with IL-25 or IL-33, normalise to sort-purified B cells from the Peyer's patches and small intestine (siLPL) (*n*=3-4 technical replicates pooled from 2-3 mice per group and representative of 2-3 independent experiments). B) Relative expression of *Slc7a8* in sort-purified lung ILC2, in comparison to bulk CD3+ T cells sort-purified from the mLN, spleen or lung (*n*=4 technical replicates pooled from 2-3 mice per group and representative of 2 independent experiments). C) Expression of *Slc7a5* and *Slc7a8* in naïve sort-purified ILC2 from the lungs of *Slc7a8*$^{+/+}$ or *Slc7a8*$^{-/-}$ mice (*n*=5 technical replicates pooled from 2-3 mice per group and representative of a single experiment).

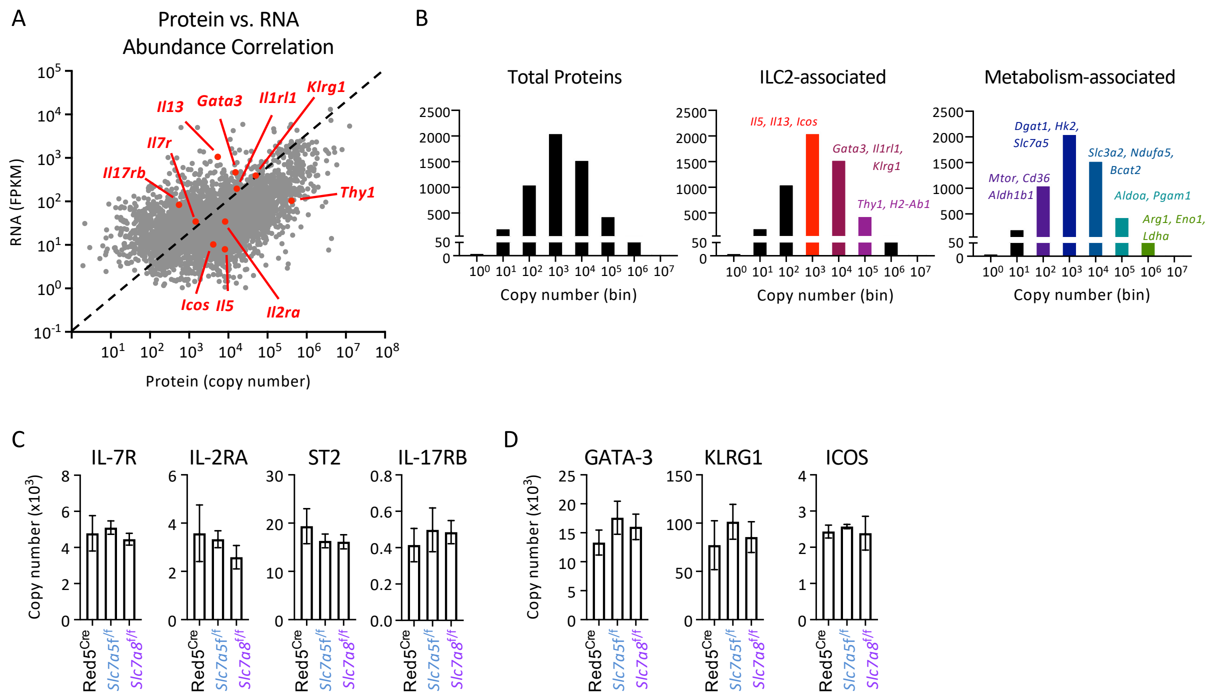


**Supplementary Figure 4. Proteomic analysis of ILC2 lacking *Slc7a5* or *Slc7a8*.** A) Comparison of relative mRNA expression and protein copy number in lung ILC2, analysed by bulk RNA sequencing and proteomics, respectively. B) Analysis of total protein copy number, and relative enrichment of key ILC2- and metabolism-associated proteins by copy number, in IL-33 elicited sort-purified lung ILC2 analysed by proteomics (*n=4* technical replicates pooled from 5 individual mice). C+D) Proteomic analysis of sort-purified lung ILC2 from IL-33 treated Red5$^{Cre}$ controls, Red5 x *Slc7a5*$^{fl/fl}$ or Red5 x *Slc7a8*$^{fl/fl}$ mice (*n*=3-4 replicates of cells pooled from 2-3 mice treated with IL-33), C) ILC and D) ILC2 associated protein copy numbers.
